# Mesenchymal stem cells therapy for spinal cord contusion: a comparative study on small and large animal models

**DOI:** 10.1101/684886

**Authors:** Yana Mukhamedshina, Iliya Shulman, Sergei Ogurcov, Alexander Kostennikov, Lena Zakirova, Elvira Akhmetzyanova, Alexander Rogozhin, Galina Masgutova, Victoria James, Ruslan Masgutov, Igor Lavrov, Albert Rizvanov

## Abstract

Here, we provided a first comparative study of the therapeutic potential of allogeneic mesenchymal stem cells derived from bone marrow (BM-MSCs), adipose tissue (AD-MSCs), and dental pulp (DP-MSCs) embedded in fibrin matrix in a small (rat) and large (pig) spinal cord injury (SCI) model during sub-acute period of spinal contusion. Results of behavioral, electrophysiological, histological assessment, as well as results of immunohistochemistry and RT-PCR analysis suggest that application of AD-MSCs combined with a fibrin matrix in a subacute period in rats (2 weeks after injury) provides significantly higher post-traumatic regeneration compared to a similar application of BM-MSCs or DP-MSCs. Within the rat model, use of AD-MSCs resulted in a marked change in (1) restoration of locomotor activity and conduction along spinal axons, (2) reduction of post-traumatic cavitation and enhancing tissue retention, and (3) modulation of microglial and astroglial activation. The effect of therapy with an autologous application of AD-MSCs was also confirmed in subacute period after spinal contusion in pigs (6 weeks after injury), however, with only partial replication of the findings observed in rats, i.e. (1) partial restoration of the somatosensory spinal pathways, (2) reduction of post-traumatic cavitation and enhancing tissue retention, and (3) modulation of astroglial activation in dorsal root entry zone. The results of this study suggest that application of AD-MSCs embedded in fibrin matrix at the site of SCI during the subacute period can facilitate regeneration of nervous tissue in rats and pigs. These results, for the first time, provide robust support for the use of AD-MSC to treat subacute SCI.

## Introduction

Mesenchymal stem cells (MSCs) have to date been one of the most promising progenitor cell types, associated with diverse functional capabilities and extensive tissue regenerative potential. MSCs can migrate to area of injury (Nitzsche et al., 2017) and integrate with damaged tissues, they also have immunomodulatory (Hashemi et al., 2013; Laroni et al., 2015; Lee & Song, 2018), anti-apoptotic, and anti-inflammatory effects due to secretion of various neurotrophic factors and cytokines (a paracrine mechanism of MSCs action), which activating signaling pathways through specific receptors on target cells (Samsonraj et al., 2017). These specific features of MSCs support their potential for treatment of neurodegenerative conditions and traumatic brain and spinal cord injury. Multiple findings suggest that MCSs could promote regeneration of nervous tissues in the brain and spinal cord (Bianco et al., 2016; Mukhamedshina et al., 2018).

MSCs with specific immunophenotypic characteristics and differentiation potential can be found in different organs. Bone marrow is the most common source of MSCs (BM-MSCs), which have been tested for spinal cord injury (SCI) therapy in preclinical and clinical studies (Sabapathy et al., 2015). There are several difficulties associated with the isolation of cells from the bone marrow, i.e. their number and regenerative potential, which decreases sharply with age. Another promising source of MSCs is adipose tissue (AD-MSCs), which due to the simple method of isolation, purification, reproduction, and proliferative activity, provides a more potential for clinical translation. AD-MSCs were found to be effective in protecting the synaptic stability of motor neurons and in the facilitation of antioxidant and anti-inflammatory mechanisms after spinal cord injury (SCI) (Ribeiro et al., 2015; Kim et al., 2015). Dental pulp-derived MSCs (DP-MSCs) have also been proposed as an alternative to BM-MSCs and AD-MSCs (Yalvac et al., 2009, 2010). DP-MSCs were found to be effective in the protection of sensory and motor neurons after SCI, through the inhibition of apoptosis, and promotion of axonal growth (Nosrat et al., 2004; Sakai et al., 2012; Yalvac et al., 2013).

The main features of all MSCs, i.e. fibroblast-like morphology, growth characteristics in culture, ability to differentiate under the influence of specific stimulants into osteogenic, adipogenic and chondrogenic lineages, expression of surface markers, and the identity of the expression profile of most genes make them valuable source for neuroregenerative studies (Minguell 2001; Wagner et al., 2005; Ryu et al., 2012; Heo et al., 2016; Wang et al., 2016). At the same time, a comparative analysis of regenerative potential of MSCs obtained from different sources indicates there are differences. BM-MSCs have greater chondrogenic and osteogenic potential with a similar colony efficiency to AD-MSCs, but lack the biological advantages of AD-MSCs when comparing proliferative capacity and immunomodulatory effects (Panepucci et al., 2004; Sakaguchi et al., 2005; Li et al., 2015). DP-MSCs, in turn, have a significantly greater cytoprotective effect compared to BM-MSCs (Song et al., 2015). We hypothesize that these differences in the features of MSCs derived from different tissue sources, as well as donor-related variations of regenerative potential of MSCs, determine the regenerative and protective effects seen when applied as a treatment for SCI.

Systemic, intraspinal and intrathecal routs of cells administration have several drawbacks, primarily associated with limited injection volumes, low stem cell viability *in situ*, and side effects. Biopolymer matrices are commonly used to maintain transplanted cells and optimize the microenvironment to increase cell survival and control distribution of bio-active molecules released by the implanted cells (Assunção-Silva et al., 2015; Caron et al., 2016; Zhao et al., 2017). The combined application of MSCs with a matrix could provide a safe and effective approach for cell transplantation in humans with SCI (Caron et al., 2016; Sabino et al., 2018; Mukhamedshina et al., 2018).

Preclinical studies of cell based therapies are commonly performed on small laboratory animals (rats, mice) with reported structural and functional restoration of damaged tissue. Nevertheless, translation of the results to clinical trials do not always produce the same effect in humans, often due to structural differences between human and small animals, as well as variation in physiological characteristics, differing mechanisms of disease development, experimental design and the inability to perform autologous transplantation (Bracken, 2009; Vandamme, 2014; Alexandrov et al., 2017). Preclinical studies performed in large animal models (for example pigs and monkeys), which share anatomical and functional similarities with humans, may facilitate better translation to human clinical trials, minimizing economic risks.

A major gap in knowledge required for the successful translation of preclinical studies is the absence of adequate comparisons between different animal models, particularly between commonly used small and large animals in the same study. To address this critical gap, we compare the therapeutic potential of allogeneic BM-MSCs, AD-MSCs, and DP-MSCs embedded in fibrin matrix (FM) in rat and pig models of spinal cord contusion. Our results indicate that in the rat model of subacute period of spinal contusion, AD-MSCs have a better neuroregenerative potential compare to other sources of MSCs, based on criteria to score functional recovery, tissue retention, reduction of post-traumatic cavitations, modulation of microglial and astroglial activation, and neuroprotection. Application of AD-MSCs in subacute period of SCI in pigs also showed significant differences based on behavioral evaluation, with some improvement in the restoration of the somatosensory pathways, tissue retention, reduction of post-traumatic cavitations, and predominant modulation of astroglial activation in dorsal root entry zone.

## RESULTS

### Distribution and survival of MSCs in the area of SCI in rats

A comparative analysis of the migration and viability of AD-MSCs, BM-MSCs and DP-MSCs transplanted into the area of SCI was conducted. In the spinal cord of rats with transplantation of transduced by LV-EGFP MSCs (3 sources - 3 groups), specific fluorescence was detected at the distance up to 5 mm rostrally and caudally from the injury site during 60 days after cells application.

At days 7 and 14, labeled EGFP AD-MSCs, BM-MSCs and DP-MSCs were predominantly located in DREZ, CC and lateral CST (Supplementary Fig.1A-F), and single fluorescent cells were present in the dorsal CST. There was no significant difference in the number and distribution of fluorescent MSCs between the experimental groups at these time points. At days 30 and 60, labeled EGFP cells were also found in the VH and VF (Supplementary Fig.1G-L). Treated animals showed tropism of transplanted MSCs to the gray matter and their preferential migration through dorsal roots of the spinal cord that can be attributed to present directional paths for their movement and the disrupted barrier in this area after SCI (Mukhamedshina et al., 2017). Collected data with distribution of MSCs in the SCI area were similar for all three experimental groups.

Whilst no significant differences were found between the experimental groups in the number of fluorescent cells in the white and gray matter in the early periods after SCI was observed, at days 30 and 60, both rostrally and caudally from the injury in the FM+DP-MSCs group, a decrease in the number of EGFP-labeled cells ∼4,1-fold (p<0.05) in VH (Supplementary Fig.1M-P) and their absence in CC compared to groups with AD-MSCs and BM-MSCs was detected.

### Behavioral results in rats

A comparative assessment of locomotor activity after application AD-MSCs, BM-MSCs and DP-MSCs was performed using the open-field Basso, Beattie, Bresnahan (BBB) (Basso et al., 1995) locomotor rating scale. On week 11 after SCI the motor function scores were higher in group with AD-MSCs [17.1±3.1] compare to other groups, with significant differences (p<0.05) within 4-11 post-injury weeks with control group FM and 4-6 weeks with BM-MSCs group (Fig.2A). During the last 4 weeks of the study, an increase (p<0.05) of the BBB scores in group with DP-MSCs application compared to FM group (Fig.2C) was detected. The lowest results on locomotor recovery among the experimental groups were found after BM-MSCs application, where the average BBB score in rats was higher (p<0.05) at 3 and 9 weeks post injury and did not change significantly on week 11 compared to control group with FM application only (Fig.2B). These findings suggest recovery of motor function with application of embedded in FM MSCs, obtained from three sources.

**Figure 1.**
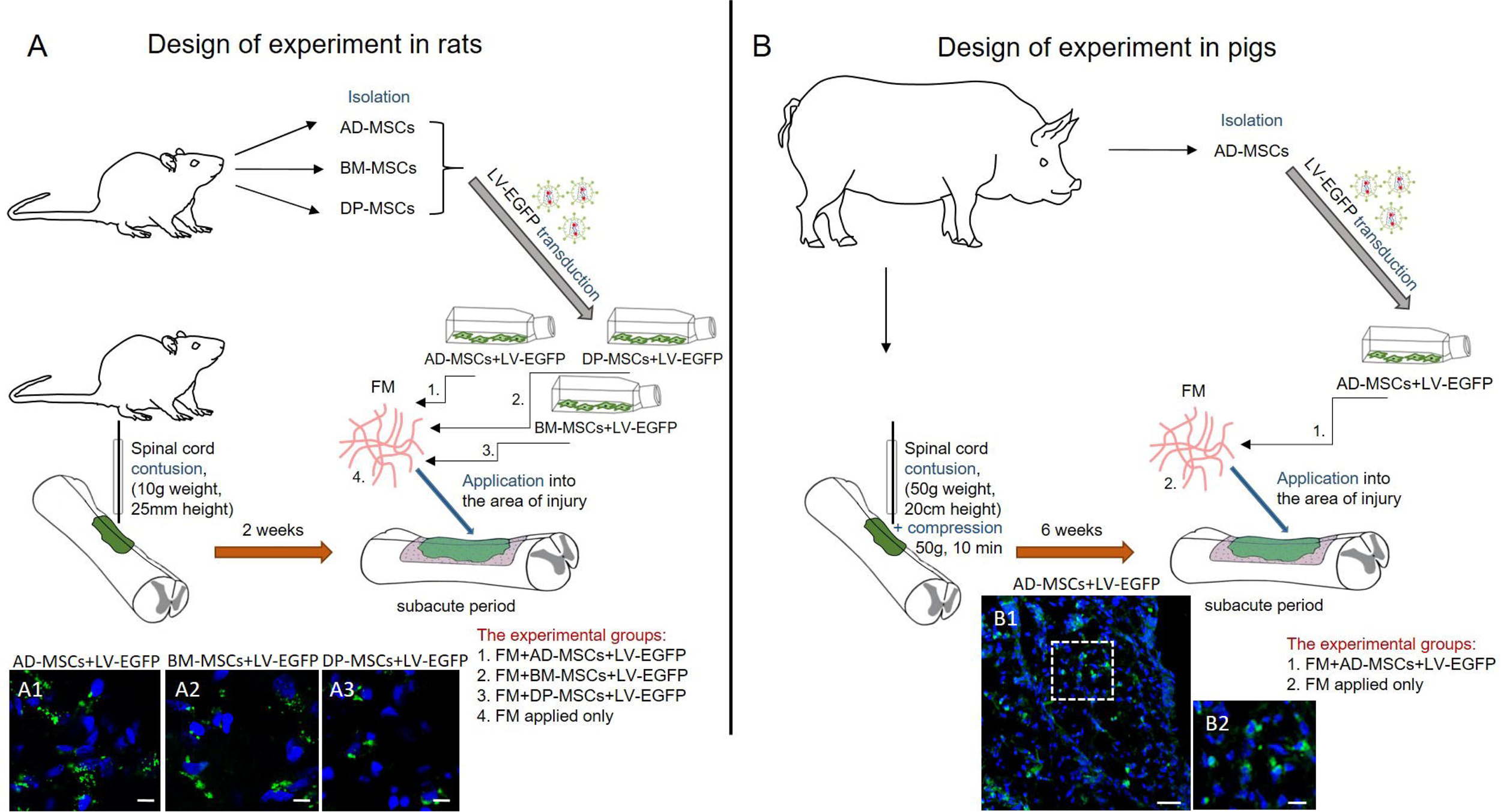
Study design in rodent and porcine models. (A) For experiments in rats AD-MSCs, BM-MSCs and DP-MSCs were isolated, transduced by LV-EGFP at passage 0 and cultivated up to passage 3. Through 2 weeks after SCI the obtained cells embedded in FM were used for allogenic application into the area of injury. (A1-A3) Below panel illustrates EGFP^+^-cells at Day 14 after application AD-MSCs, BM-MSCs and DP-MSCs, transduced by LV-EGFP, in dorsal root entry zone (DREZ). Scale bar: 5 µm. (B) For experiments in pigs AD-MSCs were isolated, transduced by LV-EGFP at passage 0 and cultivated up to passage 3. Through 6 weeks after SCI the obtained cells embedded in FM were used for autologous application into the area of injury. (B1-B2) Below panel illustrates EGFP^+^-cells at Day 14 after application AD-MSCs, transduced by LV-EGFP, in dorsal root entry zone (DREZ). The B1 area are marked with dashed boxed areas; corresponding enlarged images are shown in B2, respectively. Scale bar: 50 (B1) and 10 (B2) µm.

**Figure 2.**
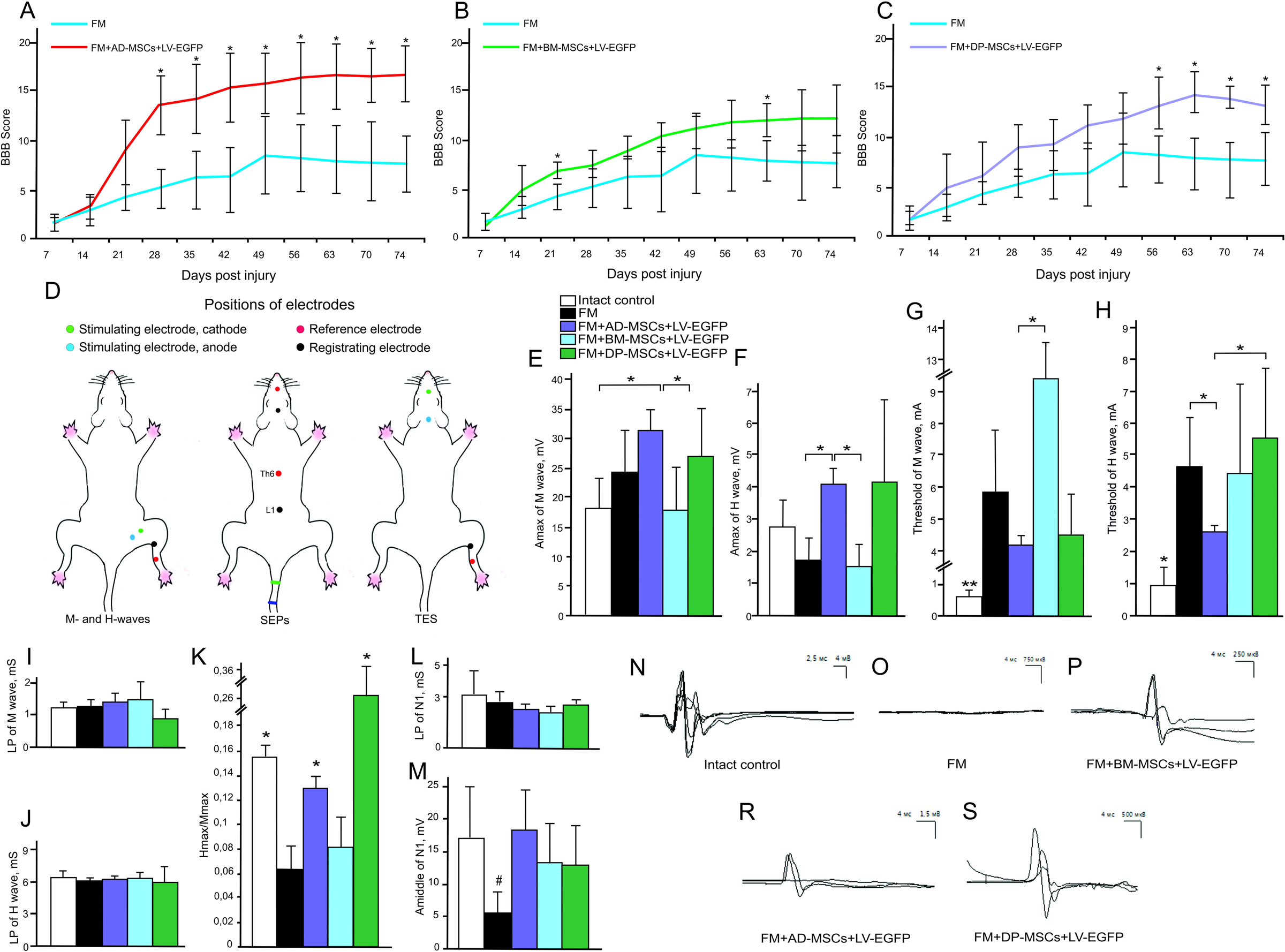
Behavioral and electrophysiological studies after SCI in experimental groups of rats. BBB locomotor scores of rats of the FM (blue line), FM+AD-MSCs+LV-EGFP (red line), FM+BM-MSCs+LV-EGFP (green line) and FM+DP-MSCs+LV-EGFP (purple line) groups (A-C). The motor function scores were significant higher after AD-MSCs and DP-MSCs application than those in the FM group for at least within 8 and 4 post-injury weeks, respectively. The lowest results in locomotor recovery between groups with cell therapy was shown after application BM-MSCs, where average BBB score was higher at 3 and 9 weeks post injury and not significantly different on week 11 compared to control group. ^□^p<0.05 as compared with the control group (FM), ^#^p<0.05 as compared with the BM-MSCs group, one-way ANOVA followed by a Tukey’s post hoc test. Schematic illustration of electrodes positions in rats (D). Amax of M- (E) and H-wave (F), threshold of M- (G) and H-waves (H), LP of M- (I) and H-waves (J), H/M wave amplitude ratio (K), LP (L) and Amiddle (M) of lumbar N1 after SCI in the experimental groups (X axis). At Days 74 after SCI the positive dynamics of the ratio Hmax/Mmax responses restoration was marked in groups with AD-MSCs and DP-MSCs application. At the same time there was a decrease of Amax of N1 in group with FM application only. □ p<0.05; □□ p<0.01; # p<0.05 as compared with intact control and experimental group with AD-MSCs application, one-way ANOVA followed by a Tukey’s post hoc test. Electrophysiology results demonstrate MEPs before SCI (N) and on week 11 in FM (O), FM+BM-MSCs+LV-EGFP (P), FM+AD-MSCs+LV-EGFP (R), and FM+DP-MSCs+LV-EGFP (S) groups.

### Electrophysiological findings

#### rats

M- and H-responses were recorded and the following parameters were evaluated across experimental groups: threshold, latency period (LP), maximum amplitude of the responses (Amax), and the ratio of H wave Amax and M wave Amax (Hmax/Mmax) (Fig.2D-K). At day 74 in group treated with AD-MSCs, a statistically significant increase of Amax (p<0.05) compared to intact control and FM+BM-MSCs treated animals was found (Fig.2E). Previously we hypothesized, that the application AD-MSCs would increase the number of muscle fibers. However, the morphometry of the calf muscles did not reveal any significant changes in the number of muscle fibers between the control and experimental groups (Mukhamedshina et al., 2018).

H-response was absent in intact rats in 33% of cases (Meinck et al., 1976) and in groups with application of AD-MSCs, BM-MSCs, and DP-MSCs in 40%, 37,5% and 25% cases, respectively. At the same time, in the control group with FM application only, the H-response was recorded in all cases, indicating facilitation of monosynaptic circuit following injury, which could be partially inhibited by supralisional influences in control and treated animals. At day 74 after SCI, the LP of H-wave remained unchanged in all investigated groups and a positive trend in the recovery (p<0.05) of the Hmax/Mmax responses rate was found in groups with AD-MSCs and DP-MSCs (Fig.2K), that could reflect the recovery from spinal shock.

In intact rats MEPs were recorded with an average amplitude 17.66±5.30 mV and latency 5.81±0.63 mc. In the experimental groups, most of the animals showed bilateral MEPs, although in 2 rats MEPs was found only on one side. The average amplitude and latency values registered by MEPs in the experimental groups presented in Table 3. In general, in groups with MSCs application a better recovery of the electrophysiological parameters was observed compared to the control group with FM application only. Comparing the ratio of the amplitude of the M-response to MEPs, all experimental groups with SCI were significantly different from the intact control, which is a sign of incomplete recovery. At the same time, the best indicators of MEPs recovery were noted in the groups with application AD-MSCs and BM-MSCs compared to other groups with SCI.

**Table 1.** Primary and secondary antibodies used in flow cytometry and immunofluorescent staining

| Antibody | Host | Dilution | Source |
| --- | --- | --- | --- |
| Thy-1 (CD90) conjugated with PE/Cy5 | Mouse | 1:100 | Biolegend |
| CD 73 | Rat | 1:100 | Biolegend |
| CD 44 conjugated with APC/Cy7 | Rat | 1:100 | Biolegend |
| CD 29 conjugated with PE | Hamster | 1:100 | Biolegend |
| CD34 | Mouse | 1:100 | Santa Cruz |
| CD45 | Rat | 1:100 | Sony |
| GFAP | Mouse | 1:200 | Santa Cruz |
| Iba1 | Goat | 1:300 | Abcam |
| Anti- goat IgG conjugated with Alexa 555 | Donkey | 1:200 | Invitrogen |
| Anti- mouse IgG conjugated with Alexa 546 | Donkey | 1:200 | Invitrogen |

**Table 3.** Results of transcranial electrical stimulation in rats

| Groups of animals | Amplitude of MEPs (mB) | Latency of MEPs (mc) | Lack of MEPs in both legs (in % of rats) | Registration of MEPs from one leg (in N. of rats) |
| --- | --- | --- | --- | --- |
| Intact control | $17.66\pm5.30$ | $5.81\pm0.63$ | - | - |
| FM | $0.5\pm0.36$ | $15.83\pm1.14$ | 44% | 1 |
| FM+AD-MSCs+LV-EGFP | $1.57\pm0.88$ | $14.08\pm3.83$ | 20% | 1 |
| FM+BM-MSCs+LV-EGFP | $1.68\pm0.7$ | $14.77\pm3.2$ | 32% | - |
| FM+DP-MSCs+LV-EGFP | $1.01\pm0.75$ | $12.85\pm3.10$ | 32% | - |

SEPs in intact rats from the lumbar level were presented by P1-N1-P2 complex, with the largest in amplitude peak N1. In intact control average values of Amiddle of peak N1 was 16.72±8.68 µV and LP – 3.08±0.97 ms (Fig.2L, M). N1-P1-N2-P2 complex (where N2 and P2 could be not registered in normal rats) was recorded with electrodes located on the scalp and Amiddle and LP values of the N1 and P1 peaks were measured as well. An average value of Amiddle the peak of N1 in intact control was 3.11±0.60 µV and LP – 7.85±0.61 ms. An average value of Amiddle the peak of P1 in intact control was 4.65±1.87 µV and LP – 14.65±1.27 mc. The complex, recorded from the lumbar level, remained intact in all groups with SCI. However, there was a significant decrease of Amiddle of N1 (p<0.05) in animals with FM application only compared to intact control and group with AD-MSCs application. The expected origin of peaks recorded from the lumbar region is the spinal roots and neurons of the posterior horns of the lumbar and sacral segments of the spinal cord. SCI at the thoracic level could affect these structures via edema and secondary circulatory changes, which could lead to a decrease in the Amiddle of the lumbar peak. The absence of these changes in group with AD-MSCs application suggests the positive effect of AD-MSCs. In contrast to the TES results, there was practically no recovery of scalp SEPs. We were able to register scalp SEPs only in one rat in the group with application AD-MSCs.

### Spinal cord morphometry in rats

Significant difference in areas of the spared tissue and abnormal cavities was found between the experimental groups (Fig.3). Application AD-MSCs lead only to improvement of morphometric characteristics at Day 60 after cell therapy compared to the control group: area of spared tissue was larger at a distance of 1 and 2 mm rostrally and 3 mm caudally from the injury epicenter (p<0.05), the total area of abnormal cavities was less at a distance 1 mm rostrally and 1-3 mm caudally from the injury epicenter (p<0.05) (Fig.3A, D, G). Experimental group treated with BM-MSCs had no significant difference from control group along 5 mm in rostral and caudal direction (Fig.3B, E, G). After application DP-MSCs, the total area of abnormal cavities was significantly less at a distance of 1 mm caudally from the injury epicenter compared to the control group (Fig.3B, E, G). At the same time, significant difference in the spared tissue was found between the groups with MSCs: 2 mm rostrally and 3 mm caudally from the injury epicenter, where this value was higher in FM+AD-MSCs group compare to FM+BM-MSCs; 2 mm rostrally and 3, 5 mm caudally from the injury epicenter, where this value is higher in FM+AD-MSCs group compare to FM+DP-MSCs. The total area of abnormal cavities in group with AD-MSCs was smaller (p<0.05) compare to BM-MSCs group in the epicenter of injury and at the distance of 3 mm caudally from this; there was no significant difference with DP-MSCs group at all tested spinal cord regions.

**Figure 3.**
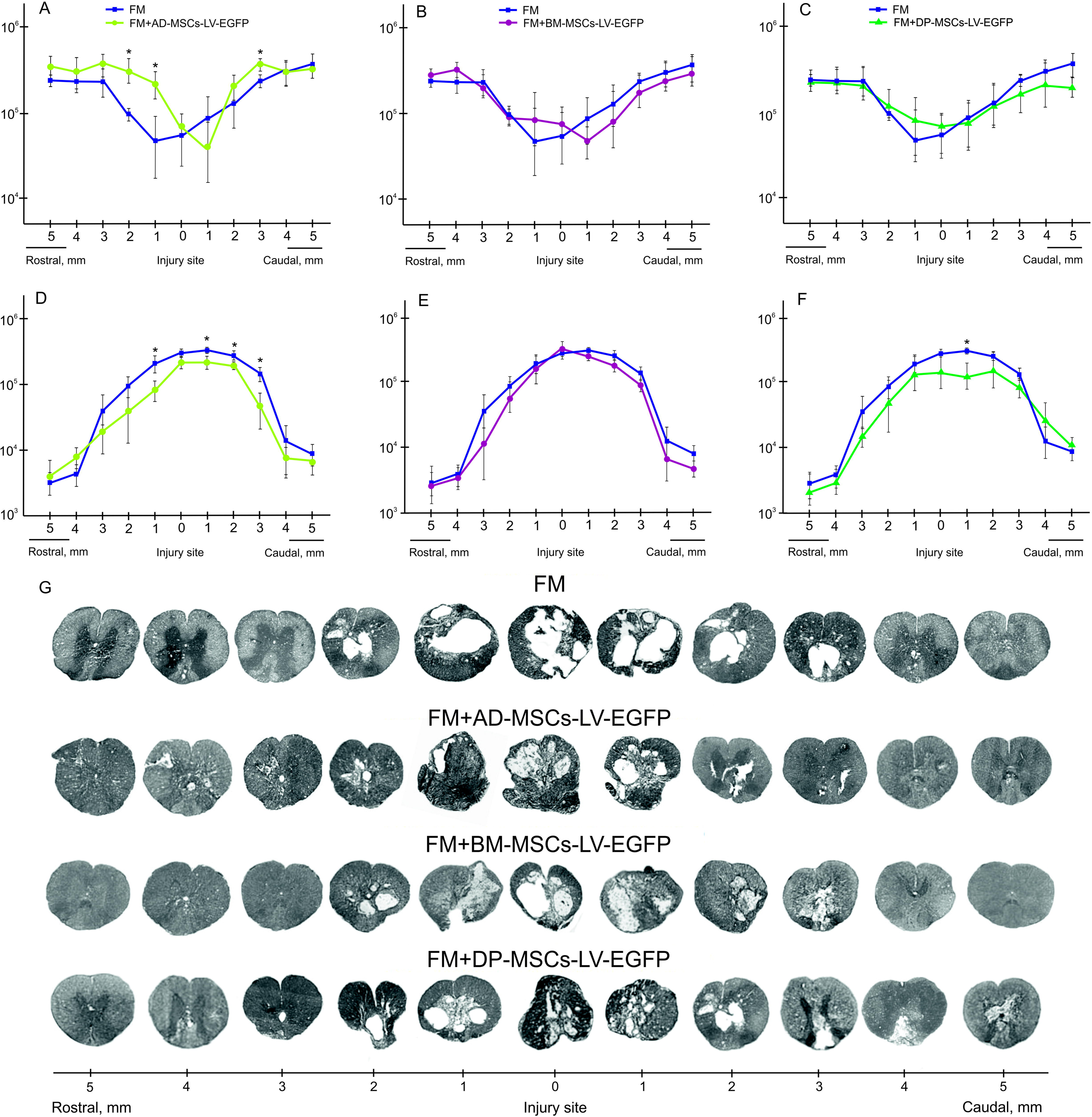
Spinal cord morphometry in experimental groups of rats. An area of the spared tissue (A-C) and a total area of abnormal cavities (D-F) 5 mm rostrally and caudally from the injury epicenter in 60 days after FM (blue line), FM+AD-MSCs+LV-EGFP (yellow line), FM+BM-MSCs+LV-EGFP (purple line) or FM+DP-MSCs+LV-EGFP (green line) application. The spared tissue in group with application AD-MSCs was higher in comparison with BM-MSCs group at the distance of 2 mm rostrally and 3 mm caudally from the injury epicenter, and in comparison with DP-MSCs group at the distance of 2 mm rostrally и 3, 5 mm caudally from the injury epicenter. The total area of abnormal cavities in group with application AD-MSCs was less in comparison with BM-MSCs group in the epicenter of injury and at the distance of 3 mm caudally from this. □ p<0.05, one-way ANOVA followed by a Tukey’s post hoc test. Cross-sections of the injured spinal cord 74 days after SCI in experimental groups at the distance of 5 mm rostrally and caudally from the injury epicenter (G). Azur-eosin staining.

### Assessment of astroglial and microglial cells in the area of SCI in rats

Using pan markers GFAP and Iba1 for astrocytes and microglia we provided a semi-quantative estimation of this cells in the areas of gray and white matter in spinal cord at a distance of 5 mm rostrally and caudally from the injury epicenter in experimental groups (Fig.4). Astroglial activation was prominent (p<0.05) in group with BM-MSCs at a distance of 5 mm rostrally from the injury epicenter in comparison with other experimental groups in VH, CC and DREZ (Fig.4A). At the same time, in rostral direction we did not find significant differences in total intensity of GFAP between FM+AD-MSCs, FM+DP-MSCs and FM application only groups, except VF for FM+DP-MSCs and FM groups (p<0.05). In the caudal direction from the injury epicenter differences between experimental groups in levels of the GFAP expression were more prominent (Fig.4B). Significant changes between groups in this direction were observed in all investigated areas except VF. However, astroglial activation was more pronounced in control group with FM application only and less in group with FM+BM-MSCs. The total intensity of GFAP was minimal in group with AD-MSCs application, where in the regions of CC and DREZ this value was less (p<0.05) compared to intact control (Fig.4B, C). Microglial activation was markedly upregulated after SCI primary at a distance of 5 mm caudal from the injury epicenter (Fig.4D). Peak values of the total intensity of Iba1 were detected in groups with FM+BM-MSCs or FM only application in grey matter - VH and DREZ. In white matter - CST and VF, the total intensity of Iba1 was higher (p<0.05) in all experimental groups compared to intact control (Fig.4D, E).

**Figure 4.**
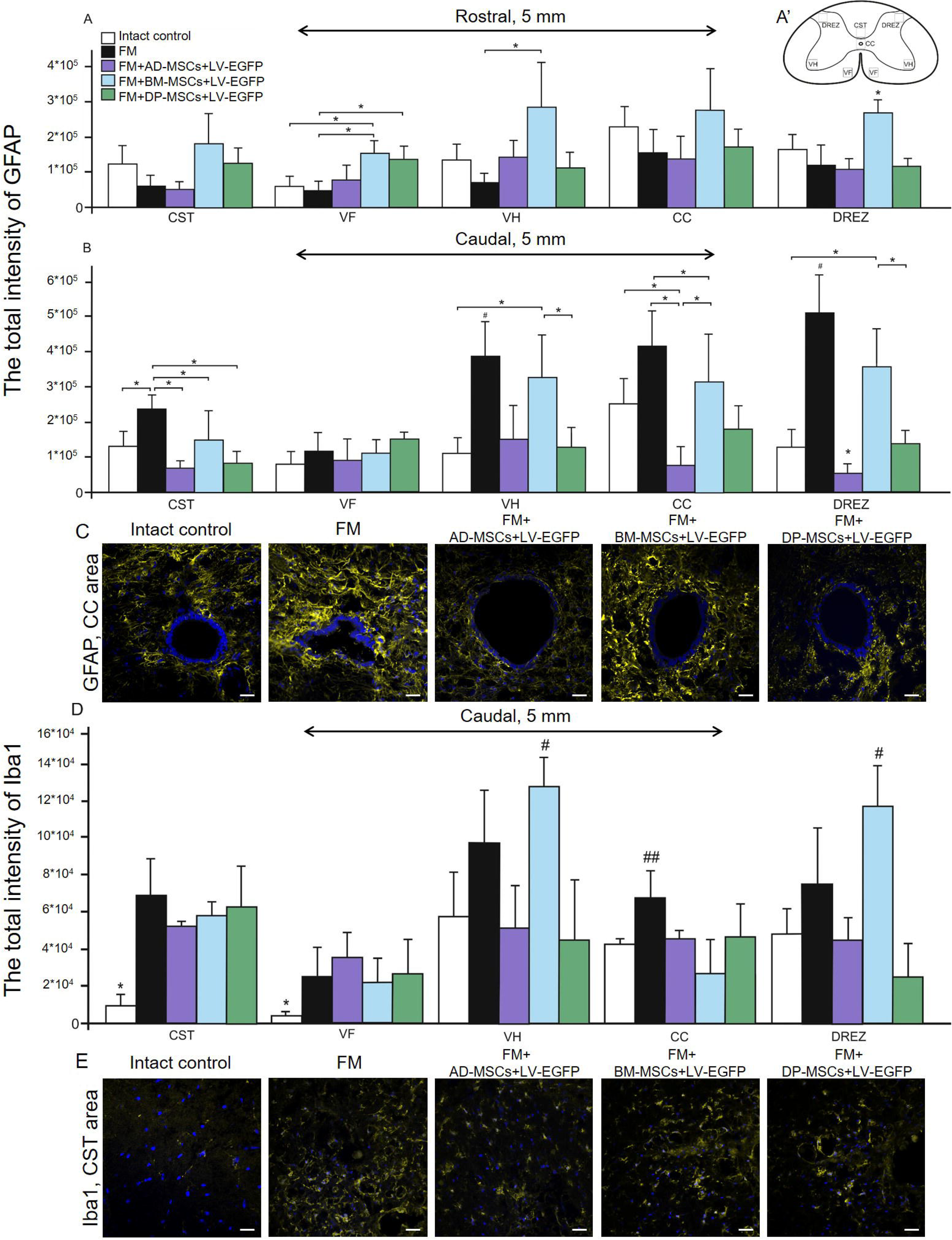
Assessment of astroglial and microglial activation at the site of injury in rats. The total intensity of GFAP (A, B) and Iba1 (D) (Y axis) in intact spinal cord (write column) or 60 days after FM (black column), FM+AD-MSCs+LV-EGFP (violet column), FM+BM-MSCs+LV-EGFP (blue column) and FM+DP-MSCs+LV-EGFP (green column) application in the examined regions. □ p<0.05; # p<0.05 and ## p<0.05 as compared with all investigated groups, except FM and DP-MSCs, respectively; one-way ANOVA followed by a Tukey’s post hoc test. (A) Schematic representation of the investigated areas: the ventral horn (VH), the main corticospinal tract (CST), the ventral funiculi (VF), the area of the central canal (CC) and the dorsal root entry zone (DREZ). (C, E) Visualization of astroglial and microglial activation using GFAP (yellow) and Iba1 (yellow) 5 mm caudally from the injury epicenter within the CC and CST in the investigated groups. Nuclei are DAPI-stained (blue). Scale bar: 25 µm.

### Analysis of mRNAs expression in the area of SCI in rats

We performed qRT-PCR on sections obtained from the injury site 74 days after SCI in experimental groups (Supplementary Fig.2). After SCI upregulation expression of neurotrophic factors mRNA such as *fgf2, vegf* and *ngf* was observed in experimental groups. Application AD-MSCs lead to the largest increase mRNA expression levels of *fgf2;* application DP-MSCs - *vegf* (p<0.05) compared to other experimental groups. Variations in expression of neural cells markers mRNAs in injured spinal cord were found. These results demonstrate that the *gfap* and *Iba1* mRNA expression was similar to those described previously by using immunohistochemical analysis. We also observed increasing mRNAs of myelin related proteins such as *MBP* and *mpz*, maximal levels that were found in group with AD-MSCs application. The mRNA expression of *pdgfβr* and *GAP-43* were markedly upregulated (p<0.05) after AD-MSCs and DP-MSCs application, when compared to another experimental groups and the intact control. Our results demonstrate that the mRNA expression of vital antiapoptotic regulator *HSPA1b* were markedly upregulated in groups with MSCs application, having the maximum value in group with FM+DP-MSCs (p<0.05).

#### Summary of results from rodent model

Results described above suggest that application of AD-MSCs combined with FM in subacute period stimulates posttraumatic regeneration to a greater extent compared to therapy with BM-MSCs and DP-MSCs. This conclusion is confirmed by (1) recovery of locomotor activity and nerve fiber conduction obtained by electrophysiological data, (2) tissue sparing and reduction of posttraumatic cavitation, (3) modulation of microglial and astroglial activation. Next we tested the effect of AD-MSCs therapy in subacute period of SCI in large animals (pigs). Approximations for the degree of injury and experimental conditions were estimated to represent those used in rodents as closely as possible to allow for comparative assessment.

### MSCs cytokine profile in pig

We provided simultaneously analyze multiple cytokine and chemokine biomarkers with Bead-Based Multiplex Assays using xMap Luminex technology, in porcine supernatant cell culture samples. Since we performed autologous transplantation in the case of experiments in pigs, we decided to analyze possible differences in the secretory phenotype of MSCs obtained from different pigs. Obtained results showed a significant difference in concentrations of cytokines/chemokines such as IFN-g, IL-6, IL-8 and IL-18 (Table 4).

**Table 4.** The AD-MSCs supernatant cytokine/chemokine concentrations (ng/ml)

| cytokine/chemokine | 1 pig | 2 pig | 3 pig | 4 pig | 5 pig |
| --- | --- | --- | --- | --- | --- |
| GM-CSF | <0.009 | <0.009 | <0.009 | <0.009 | <0.009 |
| IFN-g | <0.12 | <0.12 | <0.12 | 1.4±0.1* | 1.2±0.05* |
| IL-1a | 0.008±0 | 0.008±0 | 0.009±0 | 0.009±0.0005 | 0.008±0 |
| IL-1b | 0.01±0 | 0.01±0.0005 | 0.01±0 | 0.009±0 | 0.009±0.0005 |
| IL-1Ra | <0.03 | <0.03 | <0.03 | <0.03 | <0.03 |
| IL-2 | <0.01 | <0.01 | <0.01 | <0.01 | <0.01 |
| IL-4 | <0.05 | <0.05 | <0.05 | <0.05 | <0.05 |
| IL-6 | 2.77±0.1* | 0.15±0* | 9.21±0.3* | 0.48±0.01* | 0.57±0.05* |
| IL-8 | 21.05±0.5* | 0.21±0.01* | 14.55±0.35* | 4.56±0.2* | 3.02±0.05* |
| IL-10 | <0.009 | <0.009 | <0.009 | 0.01±0.0005 | 0.01±0.0005 |
| IL-12 | <0.03 | <0.03 | <0.03 | <0.03 | <0.03 |
| IL-18 | 0.1±0* | 0.08±0.001* | 0.09±0.001* | 0.17±0.02* | 0.35±0* |
| TNF-a | 0.01±0 | 0.01±0.0005 | 0.01±0 | 0.01±0 | 0.01±0.0005 |

\* p<0.05 as compared with analogically samples of another pigs.
< - protein concentrations below the specified value cannot be detected with the kit.

### Distribution and survival of MSCs transplanted into the area of SCI in pigs

We analyzed the behavior of MSCs 14 days after application into the site of SCI in pigs and obtained results similar to those determined in the rat model. At this time, the cells were predominantly located in the area of DREZ, dorsal horns (DH), dorsal funiculi (DF) and CC (Supplementary Fig.1Q-S). We noted that the synechias (epidural fibrosis) in pigs is more pronounced than in rats. In this regard, we also detected EGFP-labeled AD-MSCs in the area of epidural fibrosis, where more of MSCs were found above the epicenter of injury and in the caudal direction, and less - in the synechias above the rostral part of spinal cord (Supplementary Fig.1T).

### Behavioral and electrophysiology results in pigs

During the first week after SCI in pigs, no active hindlimb movements were observed, which corresponds to 1 score at the PTIBS scale (Fig.5A). At 6 weeks, before the application of FM only or FM+AD-MSCs, the animals had score 1.6±0.5 (Supplementary Video). Over the next 16 weeks after re-operation, an increase of PTIBS scores with large values in the group with AD-MSCs application was found. At the same time, no significant difference between the experimental (FM+AD-MSCs) and control (FM only) groups was found throughout the experiment, and at 22 weeks after SCI, the PTIBS scores in the groups were 4.2±0.8 and 3±1.8, respectively.

**Figure 5.**
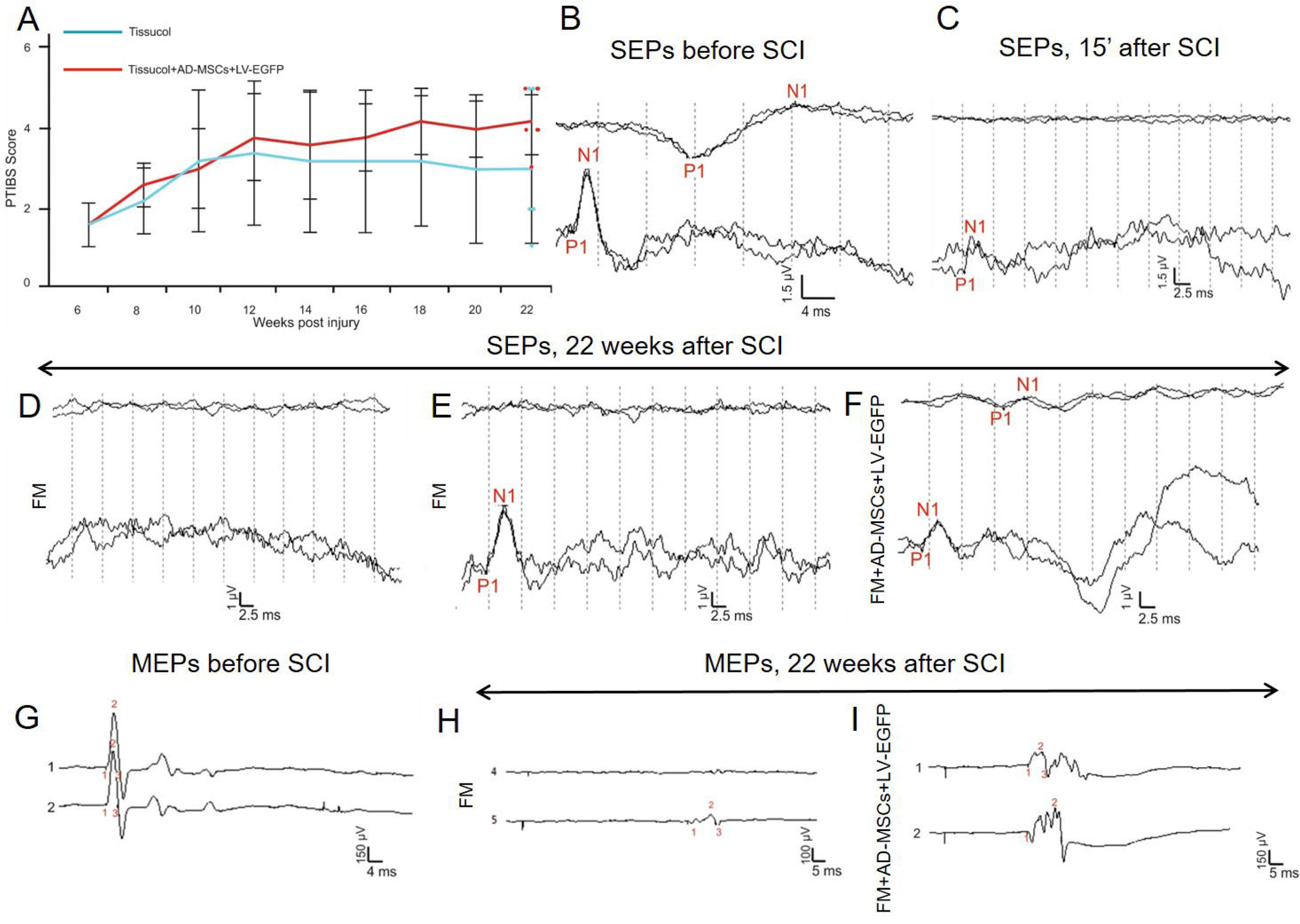
Behavioral and electrophysiology results in pigs. (A) PTIBS locomotor scores of pigs of the FM (blue line) and FM+AD-MSCs+LV-EGFP (red line) groups. Significant differences between the experimental groups throughout the experiment were not found. Electrophysiology results demonstrate SEPs (B) before SCI (reproducible peaks from lumbar and scalp electrodes) and (C) after 15 min after SCI (reproducible peaks of lesser amplitude from lumbar electrodes only). (D, E) At 22 weeks, in the control group with FM application, only 1 pig had a peak from lumbar enlargement on one side. (F) While in the experimental group with FM+AD-MSCs application, cortical peaks on one side and peaks from lumbar enlargement on both sides were recorded in 1 pigs. MEPs from the tibialis anterior muscle in pigs before SCI (G) and after 22 weeks in groups with FM (H) and AD-MSCs application (I) with registration of MEPs on one side.

Electrophysiological data (M-response, MEPs and SEPs) were recorded before the injury, at 15 minutes, 6 and 22 weeks after SCI. No significant difference in the amplitude and latency of the M-response before and after the injury was found that suggests that SCI at the thoracic level is not affecting motoneurons and motor axons of the lumbar segments. All animals before and after SCI (15 minutes) were evaluated for MEPs and SEPs (Fig.5B, C, G). At 6 weeks after SCI, no MEPs and scalp SEPs were recorded in all tested pigs. Only in one pig SEPs were recorded from lumbar electrodes. These changes showed the absence of conduction along the lateral and posterior columns of the spinal cord, which indicated the adequacy of the injury model (Skinner & Transfeldt, 2009). The absence of SEPs from the lumbar electrode in most of tested animals may indicate on the spread of spinal cord injury in caudal direction.

In experimental groups in pigs, no difference in restoration of conductivity along the lateral columns was found: in the control group with FM application after 22 weeks only one pig had MEPs on one side (Fig.5H), in the experimental group with FM+AD-MSCs application in the same period MEPs were recorded in 2 pigs on one side (Fig.5I).

The main differences were related to the ascending pathway of deep sensitivity of the spinal cord. At 22 weeks, in the control group with FM application, scalp SEPs were not found, and only one pig had a peak from lumbar enlargement on one side (Fig.5D, E). While in the experimental group with FM+AD-MSCs application, cortical peaks on one side and peaks from lumbar enlargement on both sides were recorded in one pig (Fig.5F), lumbar peaks were recorded on one side in two pigs. Thus, we can state the partial restoration of the somatosensory ways in 3 of 5 pigs with FM+AD-MSCs application.

### Spinal cord morphometry in pigs

We observed significant difference in the areas of the spared tissue and abnormal cavities between the experimental groups (Fig.6). AD-MSCs application lead to improvement of morphometric characteristics at 16 weeks after cell therapy compared to the control group (FM application only): area of spared tissue was higher at a distance of 2 mm rostrally and 1-3 mm caudally from the injury epicenter (p<0.05); the total area of abnormal cavities was less at a distance of 2 and 4 mm rostrally from the injury epicenter (p<0.05) (Fig.6A-C).

**Figure 6.**
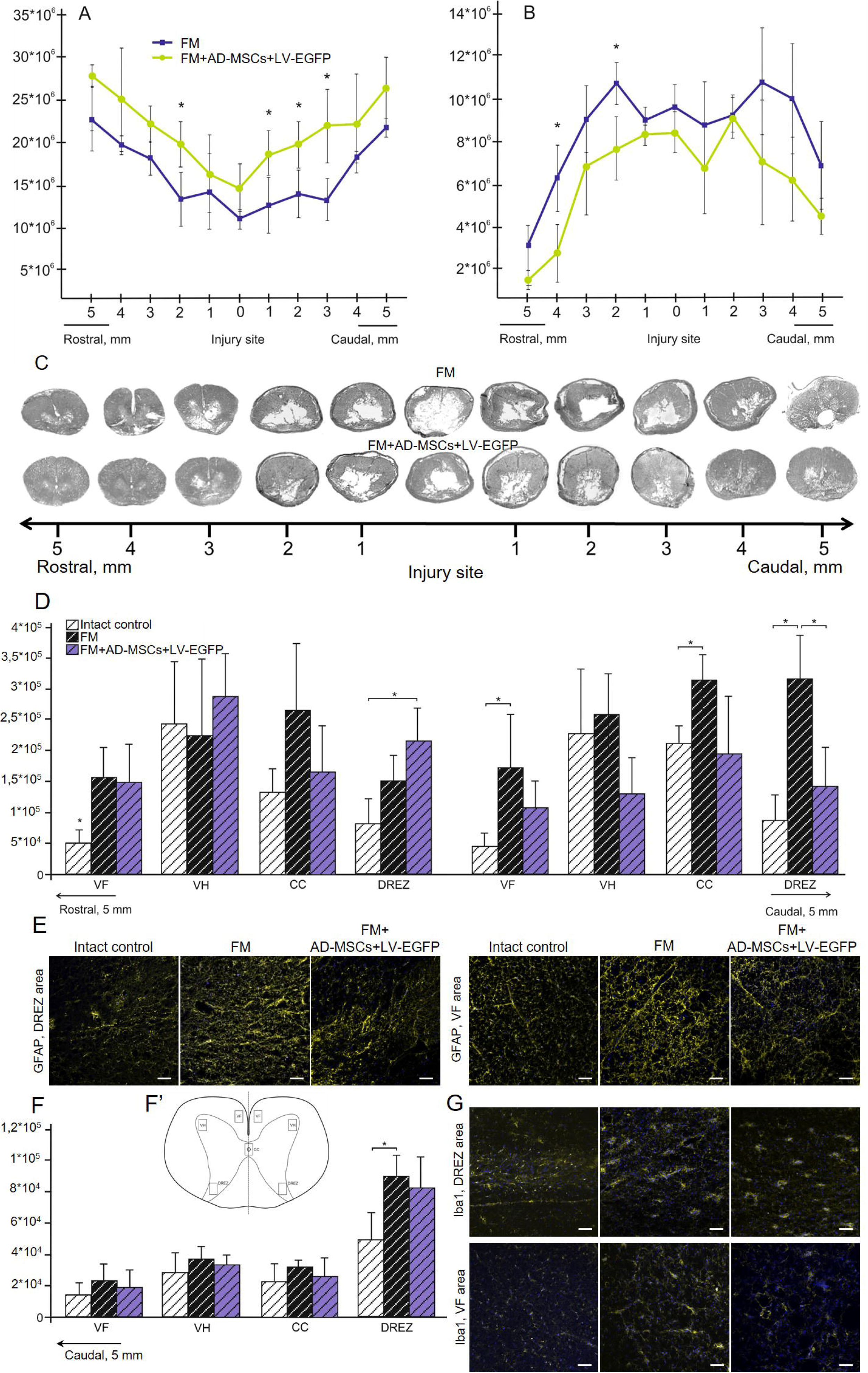
Spinal cord morphometry and assessment of astroglial/microglial activation at the site of injury in pigs. An area of the spared tissue (A) and a total area of abnormal cavities (B) 5 mm rostrally and caudally from the injury epicenter in 16 weeks after FM (blue line) and FM+AD-MSCs+LV-EGFP (yellow line) application. □ p<0.05, one-way ANOVA followed by a Tukey’s post hoc test. Cross-sections of the injured spinal cord 22 weeks after SCI in experimental groups at the distance of 5 mm rostrally and caudally from the injury epicenter (C). Azureosin staining. The total intensity of GFAP (D) and Iba1 (F) (Y axis) in intact spinal cord (write column) or 16 weeks after FM (black column) and FM+AD-MSCs+LV-EGFP (violet column) application in the examined regions. □ p<0.05 as compared with all investigated or indicated groups, one-way ANOVA followed by a Tukey’s post hoc test. (F’) Schematic representation of the investigated areas. (E, G) Visualization of astroglial and microglial activation using GFAP (yellow) and Iba1 (yellow) 5 mm caudally from the injury epicenter within the DREZ and VF in the investigated groups. Nuclei are DAPI-stained (blue). Scale bar: 50 µm.

### Assessment of astroglial and microglial cells in the area of SCI in pigs

At 22 weeks after SCI the level of GFAP expression in both pig’s experimental groups mainly increased in comparison with intact control (Fig.6D). We observed significant difference between control group with FM application only and experimental group FM+AD-MSCs in DREZ caudally from the injury epicenter, where value of the total intensity of GFAP was higher more than 2 fold (p<0.05) in control group (Fig.6D, E). After FM application, there were higher levels (p<0.05) of the GFAP expression relative to the intact controls in the VF and CC caudally from the injury epicenter, while FM+AD-MSCs group was had no significant differences in analogical areas neither with intact control, nor FM groups in pigs (Fig.10D, E).

Microglial activation was upregulated after SCI in experimental groups in DREZ at a distance of 5 mm caudal from the injury epicenter, where significant difference (p<0.05) was identified between group with FM application only and intact control (Fig.6F, G). Iba1^+^-microglia in DREZ mainly visualized as cell clusters in groups with SCI compared to intact control, in which microglia was distributed more evenly.

## DISCUSSION

Experimental approaches for the treatment of SCI have been evolved and currently include a variety of approaches with significant prevalence of genetic (Shaymardanova et al., 2013; Uchida et al., 2014; Mukhamedshina et al., 2016a) and cellular therapies. Cells of adult organism, embryonic cells, and induced pluripotent stem cells have been used in multiple studies with variable effects (Mukhamedshina et al., 2016b; Galieva et al., 2017; Manley et al., 2017; Nagoshi & Okano, 2018), as has the application of extracellular vesicles as cell free therapy (Mukhamedshina et al., 2019) and other approaches. In the context of safety and practical capacity of translation into the clinic, MSCs appear to be the most promising therapy at the current time (Xu & Yang, 2019). In addition, fibrin matrix/glue application is a widespread neurosurgical technique commonly used in humans. In this study, the comparative efficiency of MSCs derived from bone marrow, adipose tissue, and dental pulp combined with FM in subacute period of SCI was evaluated for the first time comparatively across small (rat) and large (pig) animal models.

Prior to *in vivo* experiments, we performed a morphological and phenotypic assessment of transplanted MSCs. Results showed reduced CD29 expression in BM-MSCs and DP-MSCs cultures, as well as significant reduction of CD44 expression in DP-MSCs culture in comparison with AD-MSCs. Expression of CD29 and CD44 was previously investigated in AD-MSCs, BM-MSCs, and DP-MSCs cultures depending on passage and demonstrated, that BM-MSCs show high, while DP-MSCs and AD-MSCs show low expression of CD29 at 3rd passage (Davies et al., 2015). These differences in the data between studies could reflect differences in the procedures used to isolate and cultivate MSCs. Enhanced green fluorescent protein (EGFP) has been used for a long time as a biological tag. It is considered, that EGFP gene functionally «neutral» for cell, when integrated, and provides its labeling without significant impact on basic functions (Rosochacki & Matejczyk, 2002), same as in recent study of LV-EGFP transduction on the metabolism of human placental MSCs (hPMSCs) (Yang etal., 2017). This study indicated that the expression of ectopic GFP reporter gene did not alter the metabolomics pathways in hPMSCs and did not affect their viability and phenotype profile. In our study, transduction of AD-MSCs, BM-MSCs, and DP-MSCs cells by LV-EGFP did not lead to significant changes in the expression of the investigated markers, which is consistent with other works (Yu et al., 2015).

The AD-MSCs application after SCI in rats demonstrated the best results in restoring the motor activity (BBB score). However, it would be inaccurate to estimate the effectiveness of therapy only on the basis of a behavioral test. In this regard, the summation of the data obtained from the M/H-responses, MEPs and SEPs suggests the superiority of application embedded in FM AD-MSCs compared to analogical application of FM and MSCs obtained from other sources. Results obtained with BBB assessment are often controversial between different investigators who use MSCs from different sources due to: (1) the various applied SCI models and their severity, (2) differences in transplantation protocol details (cell number, routes, and posttraumatic period of administration), (3) presence or absence of immunosuppression, (4) quality of MSCs generated in culture. At the current time, a systematic review and meta-analysis in rats models of SCI, demonstrates a difference in behavioral BBB locomotor scores means of 3.9 in favour of MSCs (Oliveri et al., 2014).

In the present study, we also found the restoration of the tissue integrity with AD-MSCs application. The reduction of a total area of abnormal cavities and improving tissue retention were reported in numerous publications on MSC transplantation in SCI (Nakajima et al., 2012; Nakano et al., 2013; Neirinckx et al., 2015; Sun et al., 2018). Ryu et al. (2012) compared the efficiency of therapy with Matrigel mixed with MSCs derived from fat, bone marrow, Wharton’s jelly, and umbilical cord blood in dogs with spinal compression and similarly to this study observed the lowest the average lesion size in AD-MSCs group. MSCs can mediate anatomical improvement in the subacute phase after SCI through anti-inflammatory activity, glial scar reducing and cell bridging effects (Qu & Zhang, 2017; Mukhamedshina et al., 2019). Our results suggest that MSCs are able to prevent the secondary injury by contributing to astroglia suppression and consistent with results of other studies with MSC transplantation in SCI (Liu et al., 2018; Ruppert et al., 2018; Krupa et al., 2018; Yang et al., 2018). We observed greater reduction of astroglial activation caudally from the injury epicenter after AD-MSCs application compared to other sources of MSCs and to control groups. At the same time, we did not find significant reduction of microglia pan markers Iba1 in the groups with MSCs transplantation.

Our findings within the rodent SCI model, encouraged us to assess the effect of AD-MSCs in subacute period of SCI in large animals. We evaluated the degree of injury and assessed the efficiency of AD-MSCs by the same criterions as in rats and we found similar results on migration MSCs after their application into the site of SCI. Our data also show tropism of transplanted AD-MSCs to the gray matter and their preferential migration through dorsal roots of the spinal cord that can be attributed to the presence of directional paths for their movement and the disrupted barrier in this area after SCI. The synechias (epidural fibrosis) was more pronounced in pigs compare to rats. In this regard, we also detected EGFP-labeled AD-MSCs in the area of epidural fibrosis, where significantly less MSCs were found in the synechias above the rostral part of spinal cord. These results may indicate the best migration of MSCs in the rostral direction from the epicenter of spinal cord injury, which confirms our earlier data on *in vitro* model (unpublished observations). At the same time, the proposed modified cell grafting procedure is non-invasive and can minimize the cell leakage.

In contrast with the rodent model, we did not find a significant improvement in motor performance and increase in behavioral scores in pigs received treatment with AD-MSCs compared to the control group (FM application only). Park et al. (2011) carry out intraspinal transplantation of UCB-MSCs in 12 hours, 1 and 2 weeks after compression SCI in dogs with significant increase in 3 different motor function scores only in the group with MSCs transplantation after 1 week. Similar therapy both in the acute (24 hours) and delayed (2 weeks) period from SCI did not lead to a motor function recovery. **S**imilar outcomes werr reported after intraspinal transplantation of Matrigel mixed with AD-MSCs, previously induced into neuronal differentiation at one week after SCI (Park et al., 2012). Electrophysiological changes we observed in pigs may suggest that FM+AD-MSCs application has a positive effect on the restoration of long spinal cord tracts. These results are consistent with other studies that show improvement in SEPs in dogs and pigs with compression SCI after UCB-MSCs and BM-MSCs intraspinal injection (Lim et al., 2007; Zurita et al., 2008). Our results also suggest that application of AD-MSCs embedded in FM affects primarily the distal to the injury site region of the spinal cord, although without significant restoration of conductivity across spinal tracts.

Similar to results in rats, application of AD-MSCs embedded in FM in SCI site in pig facilitated restoration of neural tissue integrity, as confirmed by morphometric analysis. The reduction of cavity formation and fibrosis after MSC transplantation in large animal models of SCI were reported previously (Lim et al., 2007; Zurita et al., 2008; Park et al., 2011, 2012). Assessment of astroglial activation in pigs supports the capacity of AD-MSCs to reduce GFAP expression predominantly in caudal direction from the epicenter of injury, similar to what we found in rats, although, to a lesser degree. The possibilities of reduction astroglial activation was attributed to the ability of MSCs to decrease cyclooxygenase-2, IL-6 cytokine levels (Nakajima et al., 2012; Ryu et al., 2012; Liu et al., 2018; Sun et al., 2018) and secretion of TNF-stimulated gene-6 (Wang et al., 2012; Qi et al., 2014; Song et al., 2017; Chaubey et al., 2018), which consequently decreased NF-κB signaling, modulating A1 neuroinflammatory reactive astrocytes (Lian et al., 2015; Liddelow et al., 2017). At the same time, we found no effect of AD-MSCs application on microglial activation in pigs similar to results in rats. In addition, significant differences in Iba1^+^-microglia behavior in DREZ after SCI were found in both rats and pigs. In pigs Iba1^+^-microglia in DREZ was mainly visualized as a compact cell clusters compared to pig intact control and to rats where microglia was distributed more evenly. It was reported that clusters of activated microglia are formed mainly in the areas of active demyelination of white matter near the areas of active tissue damage in the brain of patients with multiple sclerosis and this specific microglia formation was not associated with destruction of blood-brain barrier (van Horssen et al., 2012). Detected differences in microglia behavior in the area of SCI in rats and pigs not previously described and require further investigation.

### Conclusion

The results of this study demonstrate that application of AD-MSCs combined with fibrin matrix in a subacute period in rats provides significantly higher post-traumatic regeneration compared to a similar application BM-MSCs or DP-MSCs. Particularly, (1) restoration of locomotor activity and conduction along spinal axons, (2) reduction of post-traumatic cavitation and enhancing tissue retention, and (3) modulation of microglial and astroglial activation were found in rat model with AD-MSCs treatment. Besides, these data indicate that application of MSCs combined with fibrin matrix into the surface of spinal cord may be an effective and safest approach for delivering cells to the damaged area. Further, our results on pigs demonstrate partial replication of the findings observed in rats, i.e. (1) partial restoration of the somatosensory spinal pathways, (2) reduction of post-traumatic cavitation and enhancing tissue retention, and (3) modulation of astroglial activation in dorsal root entry zone.

Significant difference in functional recovery within pig groups could be attributed to the differences in the secretory phenotype of the transplanted cells (i.e autologous application in pigs in this study) and variable neuroregenerative potential of large animals. This is supported by significant differences in cytokine profile of MSCs obtained from different pigs, and previously results in donor-related heterogeneity of MSCs (Montzka et al., 2009; Boregowda et al., 2016). The therapeutic potential of adult neurogenesis is well recognized, although, the neuroregenerative capacity seems to be different between small and large animals and human that does not allows to reach an optimal clinical outcome. The application of AD-MSCs embedded in fibrin matrix at the site of SCI during the subacute period can stimulate important mechanisms of nervous tissue regeneration in both rats and pigs. Results in pigs confirm previous observations in rats and support the possible utility of AD-MSCs transplantation in large animals and in humans suffering subacute paraplegia. At the same time, we need to more carefully analyze the secretory phenotype of donor cells, confirming the therapeutic activity of transplanted MSCs with possible clinical translation.

## MATERIALS AND METHODS

The methods described herein were approved by the Kazan Federal University Animal Care and Use Committee (Permit Number 2, May 5^th^, 2015) and experimental protocols were consistent with the guidelines of the Association for Assessment and Accreditation of Laboratory Animal Care International and Physiological Section of the Russian National Committee on Bioethics. All studies were performed in such a manner as to limit the number of animals used and animal suffering. Study design with main steps illustrated in Fig.1.

### Isolation and culture of mesenchymal stem cells

For all cultured cells a medium was replaced every 3 days and at passage 3, the cells were used for the subsequent experiments. MSCs were derived from the adipose tissue, bone marrow and dental pulp of female Wistar rats (weighting 250–300 g each; Pushchino Laboratory, Russia) and adipose tissue of 4 month-old female pot-bellied pigs (9–12 kg) followed by isolation and characterization (Mukhamedshina et al., 2017a).

#### rat AD-MSCs

Rat adipose tissue was collected aseptically in an operating room. Further manipulations were carried out in a cell culture laboratory as previously described (Mukhamedshina et al., 2017a). The adipose tissue was minced, washed out by centrifugation, and digested with a 0.2% crab hepatopancreas collagenase (Biolot, Russia) at 37° C for 1 hour with shaking. Then, the homogenate was centrifuged and the enzymatic solution decanted. The cell pellet was suspended with a Dulbecco’s phosphate-buffered saline (DPBS, PanEco, Russia) solution and centrifuged to remove residual enzymes. The obtained cells were cultured in α-MEM medium with 10% fetal bovine serum (FBS), 100 U/ml penicillin, 100 μg/ml streptomycin, and 2 mM L-glutamine (all obtained from PanEco, Russia).

#### rat BM-MSCs

BM-MSCs were collected from femurs. Briefly, normal regular Wistar rats were sacrificed, and their femurs were removed with surgical scissors. After thorough and quick removal of soft tissue and cutting the epiphyses of the femurs, the marrow was collected by flushing canals with DPBS. The flushed solution was collected in 50 ml sterile tube and centrifuges 5 min at 500 g, after which the supernatant was removed. The cell pellet was suspended with in a cultural medium, containing DMEM, 10% FBS, 100 U/ml penicillin, 100 μg/ml streptomycin and 2 mM L-glutamine and thoroughly homogenized. After that, the cells were seed on the culture flasks and after 24 h, the non-adherent cells were removed by replacing the culture medium.

#### rat DP-MSCs

DP-MSCs were harvested using a previously reported method (Masgutova et al., 2017). Extracted teeth were placed in sterile PBS containing 100 U/ml penicillin and 100 mg/ml streptomycin. After the pulp was removed from the cavity, washed in sterile PBS and minced with ophthalmology scissors. After centrifugation at 2000 g for 5 min, the supernatant was discarded. Then 2,5 Mл culture medium (α-MEM, 10% FBS, 100 μM L-ascorbic acid 2-phosphate (Sigma-Aldrich, USA), 2 mM L-glutamine, 100 U/ml penicillin and 100 mg/ml streptomycin) was added to obtained cell pellet. After that, the cells were seed on the culture flasks and incubated at 37° C and 5% CO_2_ motionless for 3 days, avoiding the displacement of tissue from the bottom of plate.

#### pig MSCs

Pig adipose tissue was collected aseptically from the subcutaneous fat of 4 month-old female pot-bellied pigs under anesthesia which was induced with propofol (2-6 mg/kg) via endotracheal intubation and maintained with isoflurane (1.3%) during the whole course of the operation. Infiltration of the subcutaneous tissue was performed with a solution (total volume 200-300 ml) containing 1 ml of epinephrine hydrochloride (1 mg/ml) in 200 ml 0,9% NaCl. Small skin incision was made with a length of 0.5-1 cm. Then adipose tissue was cannulated by an endoscopic cannula (Eleps, diameter 3 mm). After that, the liquid and fat deposits are sucked off by a vacuum method. The collected subcutaneous fat was placed in a sterile 50 ml tube containing 0.9% NaCl solution and delivered to the culture laboratory for subsequent cultivation.

The obtained subcutaneous fat was centrifuged at 500 g for 5 min, after which the supernatant and pellet was removed. Preserved adipose tissue was thoroughly homogenized with sterile scissors. After, 0.9% NaCl solution was added and the centrifugation was repeated, followed by the selection of the salt solution and the precipitate. A freshly prepared sterile solution 0.2% crab hepatopancreas collagenase was added to the obtained adipose tissue homogenizate and incubated at 37° C for 1 hour with shaking. Then, the homogenate was centrifuged at 500 g for 5 min and the enzymatic solution decanted. Then the obtained cells of stromal vascular fraction of adipose tissue were seed on the culture flasks and cultivated in medium containing DMEM, 20% FBS, 2 mM L-glutamine, 100 μM L-ascorbic acid2-phosphate, 100 U/ml penicillin and 100 mg/ml streptomycin. After 24-48 hour, the non-adherent cells were removed by replacing the culture medium.

### Lentiviral transduction of MSCs

The MSCs at passage 0 were transduced with lentiviral vectors encoding enhanced green fluorescent protein (EGFP) (AD-MSCs+LV-EGFP, BM-MSCs+LV-EGFP, DP-MSCs+LV-EGFP) as previously described for subsequent transplantation in the area of SCI (Mukhamedshina et al., 2018). The percentages of EGFP-positive cells were assessed by flow cytometry (Guava EasyCyte 8HT, Millipore). After viral transduction, MSCs from different sources and animals typically begin to express EGFP within 48 hours, with a plateau reached in 96 hours and 74±5% of the cells in the population studied were EGFP-positive.

### Flow cytometry of MSCs

The cells were washed from trypsin and cultural medium in PBS by centrifuging 2 times at 500 g for 3 min. After, cells were incubated with primary antibodies (Abs) for 1 hour at room temperature (RT) (Table 1). Prior to analysis, the cells were washed and incubated with secondary Abs for 1 hour at RT. After, the cells were fixed in 10% paraformaldehyde (PFA, Sigma) for 30 min and washed in PBS by centrifuging 2 times at 500 g for 3 min. Results were evaluated using flow cytometer FACS Aria III (BD Biosciences). The obtained cells from 5 animals were analyzed in each group. Experimental data of the characterization of rat and pig MSCs are provided in Supplementary Materials.

### Animals and investigated groups

Figure 1 provides a design of experiments in rats and pigs.

#### Rats

One hundred fifteen adult female Wistar rats were randomly assigned to five groups. Allogenic MSCs+LV-EGFP, 1×10^6^ cells per rat, mixed with FM Tissucol (18 μL, Baxter) were applied on top of the injury through 2 weeks after SCI. The groups were: (1) FM+AD-MSCs+LV-EGFP, **n=25** (2) FM+BM-MSCs+LV-EGFP, **n=25** (3) FM+DP-MSCs+LV-EGFP, **n=25** (4) FM applied only, **n=25** and (5) without SCI (intact control, **n=15**). All animals were evaluated with behavioral and electrophysiological assessment. Ten animals in each experimental group and 7 in intact control were randomly selected for histology/immunohistochemistry and PCR-RT, respectively. Animals were group housed in clear plastic cages (12 h:12 h light/dark cycle) with food and water available *ad libitum.*

#### Pigs

Seventeen healthy 4 month-old female pot-bellied pigs (9–12 kg) were used in this study. All pigs were clinically judged to be in good health and to have a normal neurological status. Autologous AD-MSCs+LV-EGFP, 8×10^6^ cells per rat, mixed with FM Tissucol (150 μL) were applied on top of the injury through 6 weeks after SCI. The groups were: (1) FM+AD-MSCs+LV-EGFP, **n=5**, (2) FM applied only, **n=5** and (3) without SCI (intact control, **n=5**). Two additional pigs were used to evaluate distribution of AD-MSCs in the area of SCI at Day 14 after application.

### Surgical procedures

#### Rats

Rat SCI and cell application were carried out as previously described (Mukhamedshina et al., 2018). Rats were deeply anesthetized under general anesthesia with isoflurane and laminectomy at the Th8 vertebral level was performed. Moderate spinal cord contusion was induced with impact rod (2 mm diameter, 10 g) of a custom-made weight drop device (impactor) (Mukhamedshina et al., 2018), which centered at the Th8 and dropped from 25 mm height. After SCI the wound was sutured in layers. After surgery the rats had intramuscular injections of gentamicin (25 mg/kg, Omela, Russia) for 7 consecutive days. The injured rats’ bladders were manually emptied twice daily until spontaneous voiding occurred.

The skin was re-incised to expose the spinal cord 2 weeks after SCI (subacute period). Right after removing synechias and making several longitudinal notches in the dura mater, FM with or without MSCs were applied on top of the injury. Then the dorsal back muscles and the skin were sutured. After surgery, the rats received daily intramuscular doses of gentamicin (25 mg/kg) for 7 consecutive days. After the operation, the rats were monitored and following evaluation showed consistent and similar changes in motor performance on both legs. For cell application, we strictly selected the spinal cord contusion model that showed complete hind limb paralysis.

#### Pigs

After intramuscular injection of xylazine (0.6 mg/kg) and ketamine (5 mg/kg), anesthesia was induced with propofol (2-6 mg/kg) and maintained with isoflurane (1.3%) during the whole course of the operation. Laminectomy was performed at vertebral level Th10 and spinal cord contusion was induced by impact rod (7 mm diameter, 50 g) from 20 cm height, followed by 10 min of compression (Lee et al., 2013). Due to expected variation in spinal cord in pigs, x-ray assessment of thoracic and lumbar levels was performed to identify the same level for SCI across all animals. To approximate the degree of damage to a human SCI, sustained compression was carrying out (10 min) in addition to contusion to mimic prolong spinal compression commonly found in traumatic SCI in human (Lee, *et al*., 2013). Then, the back muscles and the skin were sutured in layers. A urinary catheter (10 Fr, Jorgensen Laboratories Inc., Loveland, CO) was manually inserted for post-injury bladder drainage and removed 3-5 days after SCI, after which the animals were able to reflexively empty their bladders. Animals were given cefazolin (25 mg/kg, Sintez, Russia) and ketoprophen (1 mg/kg, AVZ, Russia) via intramuscular injection after surgery and within 5 days, and were fasted for at least 12 hours after surgery. Pigs were housed individually for the first 48 h after which they were housed in pairs.

Under general anesthesia the skin was re-incised to expose the spinal cord 6 weeks after SCI in pigs (subacute period) that approximately corresponds to subacute period (2 weeks) in rats (Nakano et al., 2013; Kwon et al., 2015) and was also confirmed with electrophysiological findings (absence of SEPs recorded from the scalp and MEPs registered by TES) The selection of the subacute period of cell transplantation was due to the technical possibility of carrying out autologous MSCs transplantation in a clinical practice, when there are surgical indications for reoperation, during which it is possible to apply MSCs obtained from a patient in an acute period and cultivated to the required amount. After removing synechias and making several longitudinal notches in the dura mater (2-3 notches, 1-2 mm length), FM with or without MSCs were applied on top of the injured spinal cord. After the wound was sutured and pigs received intramuscular doses of cefazolin (25 mg/kg) and ketoprophen (1 mg/kg) within 5 days.

### Behavioral assessment

All behavioral studies were scored simultaneously by 2 observers who were blinded to the treatment groups. Final scores were obtained by averaging the two scores awarded by the examiners.

#### Rats

A motor function was evaluated using the open-field Basso, Beattie, Bresnahan (BBB) (Basso et al., 1995) locomotor rating scale. The baseline was obtained at three days before SCI. BBB rating scale defines three phases for the recovery of voluntary movements after spinal cord contusion: early (0 – 7 scores), intermediate (8 – 13 scores), and late (14 - 21 scores). To evaluate differences in functional recovery, a behavioral assessment in the experimental groups was performed before SCI, on day 1 and once a week up to 11 weeks after SCI.

#### Pigs

For assessment of the motor function restoration in pigs a Porcine Thoracic Injury Behavioral Scale «PTIBS» was used (Lee, *et al*., 2013). «PTIBS» is a 10-point scale, where the lowest scores «1-3» describe varying degrees of “dragging”, and the highest scores «7-10» describe walking behavior. Videotaping of locomotor recovery in the experimental groups was performed before SCI, on day 3 and once a week up to 22 weeks after SCI. All pigs were walking through a 50 m long corridor within 15 min time period, while recorded on video cameras.

### Electrophysiological studies

Electrophysiological tests were performed under anesthesia on intact rats and experimental rats at 2 and 11 weeks after SCI as previously described (Mukhamedshina et al., 2017b; Mukhamedshina et al., 2018). Electrophysiological tests were performed under anesthesia on experimental pigs before the injury, at 15 minutes, 6 and 22 weeks after SCI. The M- and H-waves (pigs only have M-response) from the tibialis anterior muscle in pigs and the gastrocnemius muscle in rats were recorded in response to stimulation of the sciatic nerve (Benavides at al., 2016; Cuellar et al., 2017).

Monopolar needle electrodes were used for recording. An active electrode was inserted into the middle of the muscle belly and the reference one was implanted within a region of the tendomuscular junction. Electrical stimulation of the sciatic nerve was performed with square-wave single stimuli and pulse width of 0.2 ms. A skin electrode with a fixed electrode spacing and monopolar needle electrodes inserted subcutaneously within an area where the sciatic nerve exits from the pelvis were used for stimulation in pigs and rats, respectively (Navarro et al., 1994).

Transcranial electrical stimulation (TES) was carried out for registration of motor evoked potentials (MEPs) recorded by subcutaneous needle electrodes in tested muscles. The recording electrodes remained in the tibialis anterior muscle or the gastrocnemius muscle. TES was performed by needle electrodes inserted under the scalp up to the contact with the bone of the skull. Cathode was placed in the middle approximately 0.5 cm caudally from the interorbital line. Anode was placed in the middle near the occipital bone. Duration of stimuli 0.04 (rats) to 0.1 (pigs) msec. with intensity from 20 to 500 V were used. An average of three repetitive and close in latency responses were analyzed and compared.

Somatosensory evoked potentials (SEPs) were registered by monopolar needle electrodes inserted subcutaneously for evaluation of conduction across the posterior columns of the spinal cord. An active electrode was inserted over upper lumbar vertebras, and reference electrode – over middle thoracic vertebras for registration from lumbar level. Cathode electrode was also inserted in the middle approximately 0.5 cm caudally from the interorbital line in rats and over the vertex in pigs, anode electrode – in the middle near the occipital bone in rats and over the snout in pigs for registration from scalp. Electrical stimulation was performed by round electrodes [please add more details about this electrode if possible: material, diameter, company name or handmade etc.] in rats. Stimulation of the tibial nerve in the medial malleolus area was performed with a cutaneous stimulating electrode with a fixed interelectrode distance in pigs. Rectangular electric stimuli 0.2 ms with frequency from 3 Hz were used. The stimulus intensity was chosen based on the tail movements in rats and foot muscles in pigs (smallest stimulus producing tail movements was used). An average of three repetitive and close in latency responses were analyzed and compared.

### Histology and immunohistochemistry

At 7, 14, 30 and 60 days after reoperation, rats were anesthetized and subjected to intracardiac perfusion with 4% PFA (4°C). Spinal cord pieces (50 mm segment centered around the injury site) were removed, postfixed in 4% PFA at 4°C overnight and incubated in 30% sucrose. At 2 and 16 weeks after reoperation, pigs were deeply anesthetized and transcardially perfused with 4% PFA (4°C). A 8 cm segment of thoracic spinal cord centered around the injury site was collected, postfixed in 4% PFA for 48 hours, divided into 1cm blocks in length and cryoprotected in 15 and 30% sucrose. Obtained rats and pigs samples embedded in tissue freezing medium (Tissue-Tek O.C.T. Compound, Sakura) and then 20 μm transverse tissue sections were obtained using the cryostat Microm HM 560 (Thermo Scientific).

A series of cross-sections obtained at 60 days in rats and 16 weeks in pigs after reoperation were used. Tissue sections prepared from spinal cord regions just 5 mm rostral and caudal to the injury epicenter was stained with Azur-eosin (MiniMed, Russia). Images were then captured with a light scanning microscope APERIOCS2 (Leica). The cross-sectional area of the spared tissue and abnormal cavities was measured. A total area of abnormal cavities in the spinal cord cross-section was calculated by adding cysts with an area of not less than 1.500 µm^2^. Aperio imagescope software was used to measure the tissue area.

For immunofluorescence staining, sections were blocked with 5% normal goat serum for 1 hour at RT and then incubated with primary Abs overnight at 4°C (Table 1). For visualization, fluorophore-conjugated secondary Abs were applied for 2 hours at RT. After washing off secondary antibodies 4′,6 diamidino-2-phenylindole (DAPI) (10μg/ml in PBS, Sigma) was used to visualize the nuclei. Coverslips were mounted on slides using mounting medium (ImmunoHistoMount, Santa Cruz) and the stained sections were examined using a LSM 780 Confocal Microscope (Carl Zeiss). Using the Zen 2012 Software (Carl Zeiss) we analyzed the total intensity of GFAP and Iba1 (semiquantitative analysis) at 5-mm increments extending from the contusion center of the SCI. All sections were imaged using identical confocal settings (laser intensity, gain and o□set). The areas selected for a semiquantitative immunohistochemical evaluation were previously described: the ventral horns (VH), the dorsal corticospinal tract (CST, except the pigs), ventral funiculi (VF), the area around the central canal (CC) and the dorsal root entry zone (DREZ) (Mukhamedshina et al., 2015). The analysis was not carried out in the CST area in pigs due to the presence of cavities in this site in control group with application FM only.

#### Real-time PCR

Total RNA was isolated from fresh rat spinal cords (5 mm long segment encompassing the injury site) in a modified phenol/chloroform extraction using Yellow Solve Kit (Silex, Russia) according to the manufacturer’s recommendations. Gene specific primers and probes were designed and the selected sets of primers/probe were blasted against the GenBank to confirm their species and gene specificity. The first strand cDNA synthesis was performed using 100 ng of total RNA, 100U of RevertAid reverse transcriptase (Thermo Fisher Scientific), 100 pmol of random hexamer primers and 5U of RNAse inhibitor according to the recommended manufacturer’s protocol. A quantitave analysis of mRNA of 18S, GFAP, Vimentin, S100, Pdgfα, Pdgfβ, Vegf, Fgf2, HSPA1b, CNPase, NGF, Iba1, mpz, Olig2, Caspase3, MBP, GAP-43 genes was performed using CFX 96 Real-Time PCR System (Bio-Rad, Hercules, CA, USA). We analyzed 100 ng of cDNA for expression of target genes, 2.5× Reaction mixture B (Syntol, Russia), 200 nM of each primer and 100 nM probe (Supplementary Table 1). The mRNA expression was normalized using 18S rRNA. Plasmid DNA with corresponding inserts was used to build a standard curve. To assess copy numbers of plasmid DNA insert we used DNA copy number calculator (http://cels.uri.edu/gsc/cndna.html). The mRNA level in intact spinal cord at Th8 level was considered as 100%. All RT-PCRs were performed in triplicate.

#### Cytokine assay

Multiplex analysis based on the xMAP Luminex technology was performed with the use of a kit for MILLIPLEX MAP Porcine Cytokine/Chemokine (magnetic) kit # PCYTMG-23K-13PX (Merck), according to the manufacturer’s instructions. Experiments were performed in triplicate. The kit enables a simultaneous multiplex analysis of 13 cytokines/chemokines/interleukins pig (GM-CSF, IFN-g, IL-1a, IL-1b, IL-1Ra, IL-2, IL-4, IL-6, IL-8, IL-10, IL-12, IL-18, TNF-a) in a 25-µl porcine supernatant cell culture samples at passages 3.

#### Statistical analysis

All statistics were calculated using Origin 7.0 SR0 Software (OriginLab, Northampton, MA, United States). Data are presented as mean ± standard deviation (SD). To determine statistical significance of the behavioral, electrophysiological, and morphometric data between experimental groups, which included different numbers of animals, we used nonparametric methods for testing whether samples originate from the same distribution. A one-way analysis of variance (ANOVA) with Tukey’s test or two-way analysis of variance (ANOVA) were used in this study for multiple comparisons between tested groups. All analyses were performed in a blinded manner with respect to the treatment group. A value of p<0.05 was considered statistically significant.

## Acknowledgments

We thank EE Garanina (Kazan Federal University, Kazan, Russia) for assistance in some of the experiments. The study was supported by grants 16-34-60101 (Y.O. Mukhamedshina) from Russian Foundation for Basic Research. This work was performed in accordance with the Program of Competitive Growth of the Kazan Federal University. AA Rizvanov was supported by state assignment 20.5175.2017/6.7 of the Ministry of Education and Science of Russian Federation.

---

**Supplemental Fig. 1. Distribution and survival of MSCs encapsulated in fibrin matrix after application into the area of SCI in rats (red box) and pigs (blue box).** MSCs predominantly migrate through the posterior roots of the spinal cord. In the early periods (7-14 days post-transplantation) labeled by LV-EGFP (green) AD-MSCs, BM-MSCs and DP-MSCs were located mainly in dorsal root entry zone (DREZ, A-C) И central canal (CC, D-F) in rats. On late periods (30-60 days post-transplantation) labeled by LV-EGFP cells were observed also in the area ventral horns (VH, G-I) И ventral funiculi (VF, J-L) in rats. The A-L areas are marked with dashed boxed areas; corresponding enlarged images are shown in A’-L’, respectively. At Day 14 after application labeled by LV-EGFP MSCs (green) mainly located in the area of dorsal root entry zone (DREZ), dorsal funiculi (DF, Q), dorsal horns (DH, R) and central canal (CC, S) in pigs. The Q and R areas are marked with dashed boxed areas; corresponding enlarged images are shown in Q’ and R’, respectively. A scheme of MSCs distribution in the area of epidural fibrosis (gray line above spinal cord longitudinal section) according to the distance from injury site (T). MSCs were also detected in the area of epidural fibrosis, where more of MSCs were found above the epicenter of injury and in the caudal direction, and less - in the synechias above the rostral part of spinal cord. Asterisk indicate CC. Color dashed boxes (brown – rostral part, blue – epicenter of injury, green – caudal part) in scheme indicate corresponding area with MSCs distribution. Nuclei are stained with DAPI (blue). Scale bar: 20 (A-L), 10 (H’) and 5 μm (A’-G’, I’, J’-L’). 50 (Q, R), 20 (S) and 10 μm (Q’, R’, T’, E’, brown and green dashed boxes).

**Supplemental Fig.2. Analysis of mRNAs expression in the area of SCI in rats.** *Gfap, fgf2, HSPA1b, CNPase, pdgfβr, S100, vegf, ngf, olig2, caspase 3, MBP, GAP-43, pdgfαr, iba1, mpz* and *vimentin* mRNA expression in intact spinal cord (write column) and in 60 days after FM (black column), FM+AD-MSCs+LV-EGFP (purple column), FM+BM-MSCs+LV-EGFP (blue column) or FM+DP-MSCs+LV-EGFP (green column) application. The mRNA expression levels in intact spinal cord was considered as 100%. □ p< 0.05; # p<0.05 as compared with all investigated groups, except DP-MSCs; ## p<0.05 as compared with all investigated groups, except AD-MSCs, one-way ANOVA followed by a Tukey’s post hoc test.

