## Supplementary Information for "Mesenchymal stem cells therapy for spinal cord contusion: a comparative study on small and large animal models"

Mukhamedshina et al.  **Supplemental Material 1**

**Characterization of rat mesenchymal stem cells**

The analysis of the surface markers expression in primary cultures of AD-MSCs, BM-MSCs and DP-MSCs, native and transduced with LV-EGFP, by flow cytometry were performed (Supplementary Таble 1). The expression of CD34 and CD45 was below the threshold <2% in all obtained cell cultures. The expression of CD90 and CD73 is considered to be predominant and more than 90-95% of cells in the MSC population should express these surface markers (Dominici, et al. 2006). The highest rates of CD90 and CD73 were observed in AD-MSCs 99.9±0.17/98.6±2.3% and 94±0.5/87.5±4.4%, respectively. The level of CD90 expression was more than 90% for all tested cultures, however, a significantly lower (p<0.05) value was found in native and transduced by LV-EGFP DP-MSCs compared to AD-MSCs and BM-MSCs. BM-MSCs and DP-MSCs cultures had reduced expression of CD29 (p<0.05) compared to AD-MSCs. CD44 expression was significantly reduced (p<0.05) in cultures both native and transduced LV-EGFP DP-MSCs compared to BM-MSCs and AD-MSCs. It should be noted that transduction by LV-EGFP AD-MSCs, BM-MSCs, and DP-MSCs did not lead to significant changes in the expression of the tested markers.

**Supplementary Таble 1.** Flow cytometry data on the expression of surface antigens in cells derived from rats adipose tissue, bone marrow and tooth pulp before/after transduction by LV-EGFP (%)

| **Source of MSC** | **Thy-1 (CD90)** | **CD73** | **CD44** | **CD29** |
| --- | --- | --- | --- | --- |
| Adipose tissue | 99.9±0.17/98.6±2.3 | 94±0.5/87.5±4.4 | 98.4±2.9/98.7±2.6 | 88±4/80±8.5 |
| Bone marrow | 98.9±1.6/96.8±5.8 | 91±0.5/88±4 | 90.6±5.8/93±11.3 | 59±7.3/51±9* |
| Dental pulp | 91±0.5/88.5±1.2*^#^ | 87.7±1.7/87±8 | 74±0.5/69±9%*^#^ | 49±5.3/44±0,5* |

* p<0.05 as compared with analogically samples of AD-MSCs cultures, ^#^ p<0.05 as compared with analogically samples of BM-MSCs cultures

**Supplemental Material 2**

**Characterization of pig mesenchymal stem cells obtained from adipose tissue**

The adherent cells in the primary culture were located single or in small groups at Day 1-2 after seeding of the cells derived from adipose tissue. On the 2 day of cultivation, the cells began to divide, forming islets. All cells were similar morphologically, had a small size and an elongated (fibroblast-like) shape, in most cases with several processes. With further cultivation, the cells spread over the bottom of a culture flask and increased in size, preserving the fibroblast-like morphology. As the number of cells in a culture increased, their processes more closely adjoined each other and the borders of individual cells were difficult to distinguish. Indicated morphology that typical for MSC, cultured cells were maintained for at least 6 passages.

Based on the flow cytometry (before and after transduction), data that indicating the expression of MSC markers by the obtained cells were received: Thy-1 (CD90) – 94.5±3.7/93.6±5.2%, CD73 - 92±2.5/91±0.5%, CD44 – 86.7±8.9/78±4% and CD29 - 63±9.5/52.6±13.5%. However, the expression of such surface antigens as CD34 and CD45 in AD-MSCs was not detected.

**Supplemental Table.** Primers and probes for RT-PCR

| **Primer** | **Nucleotide sequence** |
| --- | --- |
| 18S-TM-Forward | gCCgCTAgAggTgAAATTCTTg |
| 18S-TM-Reverse | CATTCTTggCAAATgCTTTCg |
| 18S-TM-Probe | [HEX]ACCgCgCAAgACggACCAg [BH2] |
| V164-TM-Forward | TATATCTTCAAgCCgTCCTgTg |
| V164-TM-Reverse | TCTCCTATgTgCTggCTTTg |
| V164-TM-Probe | [FAM]TCCgCATgATCTgCATAgTgACgTTg [BH2] |
| Vim-TM-Forward | ACCCTgCAgTCATTCAgACA |
| Vim-TM-Reverse | TCCTggATCTCTTCATCgTg |
| Vim-TM-Probe | [HEX] CTggCACgTCTTgACCTTgAACg [BH2] |
| S100b-TM-Forward | GAgAgAgggTgACAAgCACA |
| S100b-TM-Reverse | CACCACTTCCTgCTCTTTgA |
| S100b-TM-Probe | [FAM] CgAgCTCTCTCACTTCCTggAggAA [BH1] |
| GFAP-TM-Forward | TTTCTCCAACCTCCAgATCC |
| GFAP-TM-Reverse | CTCCTTAATgACCTCgCCAT |
| GFAP-TM-Probe | [FAM] CCgCATCTCCACCgTCTTTACCA [BH1] |
| PDGFRa-TM-Forward | GgTTAgAggAgCACCTggAg |
| PDGFRa-TM-Reverse | TCTCACCTCACATCCgTCTC |
| PDGFRa -TM-Probe | [FAM] ATgCgCgACCTCCAACCTgA [BH1] |
| PDGFb-TM-Forward | CTgCAATAACCgCAATTgTg |
| PDGFb-TM-Reverse | TCgATCTTTCTCACCTgCAC |
| PDGFb-TM-Probe | [FAM] CCgCATCTgCACCTgCgAg [BH1] |
| FGF2-TM-Forward | GCTgCTggCTTCTAAgTgTg |
| FGF2-TM-Reverse | GTgCCACATACCAACTggAg |
| FGF2-TM-Probe | [FAM] TCTTCTTTgAACgCCTggAgTCCA [BH1] |
| HSPA1b-TM-Forward | CCAGGCAGGACCCAATCACA |
| HSPA1b-TM-Reverse | CGCAAGGTAGCGGTCTCTGT |
| HSPA1b-TM-Probe | (6-FAM)–CCGCCCAGCACTTTCAGGAGCTGACCC [BH1] |
| CNPase-TM-Forward | AGACATAGTGCCCGCAAAG |
| CNPase-TM-Reverse | GCTTGTCCTTAGCTCCTGAG |
| CNPase-TM-Probe | (6-FAM)–AGCCACACATTCCTGCCCAAGAT [BH1] |
| NGF-TM-Forward | CCAAGGACGCAGCTTTCTAT |
| NGF-TM-Reverse | CTCCGGTGAGTCCTGTTGAA |
| Iba1-TM-Forward | ACCAgCgTCTgAggAgCTAT |
| Iba1-TM-Reverse | AggAAgTgCTTgTTgATCCC |
| Iba1-TM-Probe | [HEX] CCCTgCAAATCCTTgCTCTggC [BH2] |
| mpz-TM-Forward | TCgCAAAgATgAgCAgAg |
| mpz-TM-Reverse | ggCCCATCATgTTCTTgA |
| mpz-TM-Probe | [FAM]CCAgTAGAACCAgCCTCAAgAAC [BH1] |
| Olig2-TM-Forward | AGTGCGCGATGCTAAGCTCT |
| Olig2-TM-Reverse | TGGGCCACGACACAGAAAGA |
| Olig2-TM-Probe | (6-FAM)–CGCGCCTCGTCGTCTAAGCCCGC [BH1] |
| Caspase3-TM-Forward | AATTCAAGGGACGGGTCATG |
| Caspase3-TM-Reverse | GCTTGTGCGCGTACAGTTTC |
| Caspase3-TM-Probe | [HEX] CggCCTCCACTggTATTTTATgACACg [BH2] |
| MBP-TM-Forward | ACACGGGCATCCTTGACTCCATCGG |
| MBP-TM-Reverse | TCCGGAACCAGGTGGTTTTCAGCG |
| GAP-43-TM-Forward | GCAGAAAAGAGGTGGAGAGG |
| GAP-43-TM-Reverse | TTGTTCAATCTTTTGGTCCTCATC |
| GAP-43-TM-Probe | [FAM] AGAGAAGGCAGGAAGAAGGCAGG [BH1] |
