## Supplementary figures and images for "Mesenchymal stem cells therapy for spinal cord contusion: a comparative study on small and large animal models"

### Supplementary figure 1

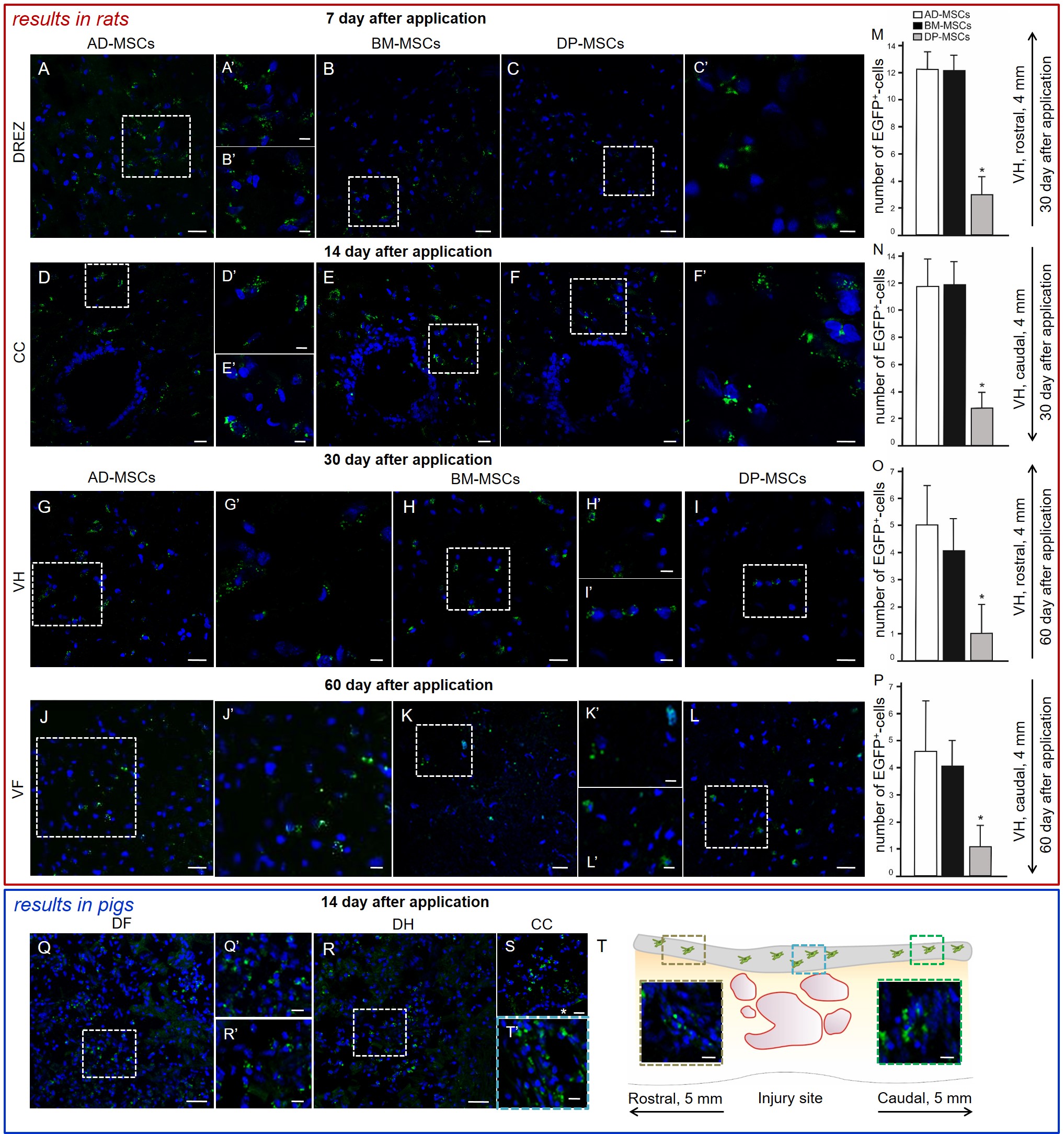

### Supplementary figure 2

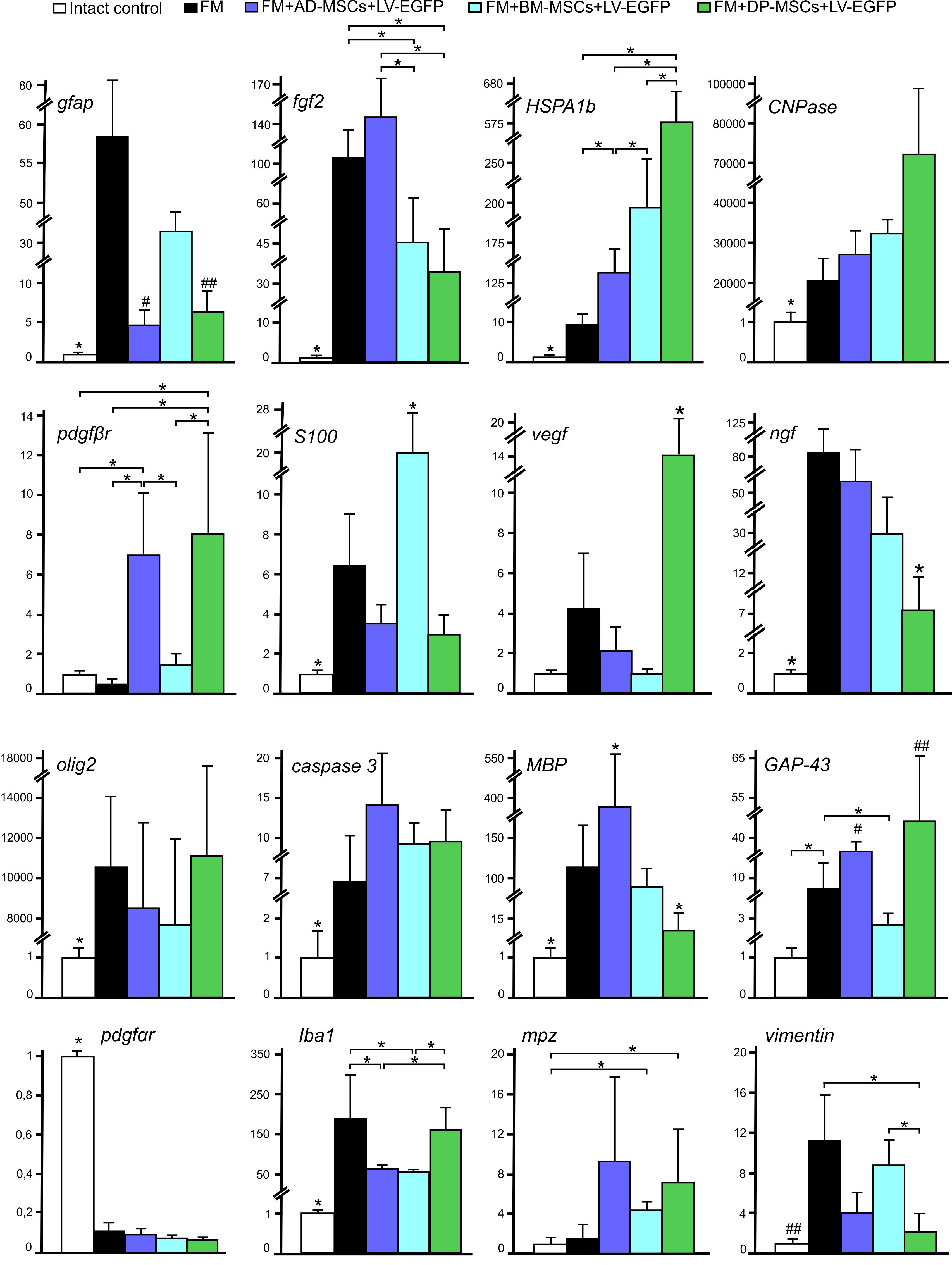
